# Brainways: An AI-based Tool for Automated Registration, Quantification and Generation of Brain-wide Activity Networks Based on Fluorescence in Coronal Slices

**DOI:** 10.1101/2023.05.25.542252

**Authors:** Ben Kantor, Inbal Ben-Ami Bartal

## Abstract

A central current trend in neuroscience involves the identification of brain-wide neural circuits associated with complex behavior. A major challenge for this approach involves the laborious process for registration and quantification of fluorescence on histological brain slices, as well as the difficulty of deriving functional insight from the complex resulting datasets. As a solution, we developed Brainways, a simple-to-use AI-based open-source software for the identification of neural networks involved in a specific behavior, from digital images to network analysis. Brainways offers automatic registration of coronal slices to any 3D brain atlas, and provides quantification of fluorescent markers (e.g. activity marker, tracer) per region, as well as statistical comparisons with visual mapping of contrasts between conditions. A built-in partial least squares task analysis provides the neural patterns associated with a specific contrast, as well as network graph analysis representing functional connectivity. Trained on atlases for rats and mice, Brainways currently provides above 80% atlas registration accuracy and allows the user to easily adjust the outputs for better fit. Below, a case study validation of Brainways is demonstrated on a previously published data set describing the neural correlates of empathic helping behavior in rats. The original results were successfully replicated and expanded upon, due to the exponentially larger sample size that covered over a 100 times more brain tissue compared to the original manual sampling. Brainways thus provides a fast, accurate solution for quantification of large-scale projects and facilitates novel neurobiological insights about the structural and functional neural networks involved in complex behavior. Brainways has a highly accessible GUI and is functionality exposed through a Python-based API, which can be enhanced for different applications.

## Introduction

Fluorescent tagging for immediate early genes (IEG) has been increasingly used to identify functional networks that participate in complex behaviors (Vetere et al., 2017; Wheeler et al., 2013; Renier et al., 2016; Turkheimer et al., 2022). While this strategy is not without caveats, such as low temporal resolution and the use of an indirect index for activity, it has led to important insights and is being globally adopted. The advantage of this approach is that it provides unbiased data-driven hypotheses for follow-up manipulations on specific neural projections and subpopulations. Furthermore, activity tagging can be combined with other markers of cell identity, such as cell type, receptors, or a tracer, to provide enhanced activity-identity mapping of subpopulations of interest.

While this approach shows promise, several obstacles have impeded progress. First, quantification and analysis are difficult, especially for high-resolution datasets. Analyzing fluorescence in histological images requires registration to a common reference brain atlas, followed by quantification of fluorescence levels or number of marker-positive cells in each brain region of interest (ROI). Due to a lack of available user-friendly tools to perform registration and quantification in a fast, streamlined, and accurate manner, this process is often performed manually, limiting the scope of the investigation to specific ROIs or a low sampling rate in order to reduce the quantification timeline. Unlike manual analysis, which requires expertise, is labor-intensive and prone to inter-rater variability, automatic registration and quantification provides efficient and reliable analysis of entire digital sections. Another substantial challenge lies in analysis and interpretation of brain-wide quantification, especially as the bar continues to rise in terms of resolution and cell specificity. Tools that use brainwide activity mapping data to generate neural network “connectomes” are needed to bridge this gap.

To this end, we developed Brainways, an open source, rapid, and user-friendly AI-based software that provides an automated pipeline for the analysis of fluorescence tagging on coronal sections. Brainways aims to address these challenges and provide a platform for collaboration between labs using different model organisms for studying the brain. Brainways provides an easy way to perform automated registration, quantification, and statistical analysis of histological brain slices from a whole experiment. Brainways provides an easy-to-use graphical user interface (GUI) that allows for adjustment of the automatic registration if needed. Brainways performs cell detection and quantification, maps detected cells to registered ROIs, provides positive cell counts per region, and performs statistical comparisons between subjects in different experimental conditions, providing a short turn-around from slide scanning to brain-wide quantification and analysis.

Brainways GUI provides a graphical fMRI-like visualization of the neural pattern associated with a specific contrast, as well as the network graph analysis based on interregion correlations. The registration is performed using a deep learning algorithm trained on data that was annotated using the Brainways GUI. Coronal slices are mapped on the anterior-posterior axis and the visible hemisphere is detected (left, right, or both). Nonrigid registration is performed using Elastix (Klein et al., 2010), and cell detection is performed using StarDist (Schmidt et al., 2018).

Brainways presents enhanced functionality compared to existing software solutions. Several algorithms exist for histological slice registration (Carey et al., 2023; Song et al., 2020; Xiong et al., 2018), which are only suitable for mouse brains. Additionally, all of these algorithms require integration with other software to complete the analysis, resulting in a cumbersome workflow and often requires programming experience. Brainways offers functional insights on the brain-wide activity of a wide range of behaviors of interest across species, accessible to users at any level working with coronal slices, used by many labs for their high accessibility and cost-effectiveness.

In order to examine the validity and reliability of the quantification provided by Brainways, we re-analyzed a manually quantified data set previously published by our group (Ben-Ami Bartal et al., 2021). In this use case, the main findings were reliably replicated despite the difference in quantification methods and the much greater coverage of neural tissue used in the reanalysis compared to the original quantification. The data set consists of neural activity indexed via the IEG c-Fos of rats tested in the Helping Behavior Test (HBT). The study aimed to outline the neural activity associated with prosocial motivation to release a trapped conspecific. Brainways-based analysis replicated the original findings, providing an enhanced neural network for prosocial motivation. Moreover, the Brainways-assisted reanalysis took a fraction of the time of the manual quantification, which had required multiple experimenters working over many months. In summary, Brainways is a unique tool that can lead to the discovery of novel neurobiological mechanisms associated with complex behavior. The code is freely available on GitHub^1^, along with an instructional video demonstration.

## Results

Brainways is composed of five modules that combine to lead from the digital image to an excel output of positive cell numbers from a whole cohort of subjects (Fig. 2). These modules can be accessed either through the Python API or through a GUI for easy access for users who lack coding skills. The “Atlas Registration” module takes a slice image and registers it to the selected brain atlas of choice. The Waxholm Space Rat Atlas (Papp et al., 2014; Osen et al., 2019) and Allen mouse brain atlas (Wang et al., 2020) were used for all data presented in this article. The “Rigid Registration” module takes the slice that was registered using the previous module, computationally separates the tissue from the background, and performs an affine transformation to match the input tissue to the atlas slice at the registered location. The “Non-Rigid Registration” module then takes the output of the previous module and performs an elastic registration using the Elastix Thin-Plate-Spline interpolation algorithm. The output of all previous modules can be visualized and manually modified using the Brainways GUI. The “Cell Detection” module is used to perform cell/nucleus detection from within Brainways using the StarDist algorithm (Schmidt et al., 2018), or by importing cell detections to Brainways from another software (e.g. QuPath). The detected cell locations are then registered to the atlas by applying the previous registration steps to the cell locations. Finally, the “Analysis” module takes the result of all previous modules, outputs normalized cell counts for each brain region, and allows further statistical analysis, such as ANOVA contrast analysis and network graph-based analysis.

### Atlas Registration

The purpose of the Atlas Registration module is to find the 3D location of a coronal rat brain slice image in the reference atlas (Fig. 2A). The Atlas Registration module receives a coronal rat brain slice image and outputs the slice’s location in the Anterior-Posterior (AP) axis of the atlas, rotation of the slice relative to the atlas in three rotation axes, and which of the hemispheres is visible in the slice. Because the rough features of the slice are visible in low resolution, the input images are downscaled and fed to a deep neural network that was trained to output the AP axis and the visible hemisphere classification. The registration parameters, including the 3D rotation parameters, can be manually adjusted using sliders in the Brainways GUI. The Atlas registration algorithm can also be accessed using the Python API from any Python script.

The Brainways GUI can be used to register image slices for a variety of atlases from different species. This is achieved by using the Brainglobe Atlas API (Claudi et al., 2020) for atlas retrieval and access in Brainways. Since there was no rat atlas available in the Brainglobe repository, we adjusted and added the Waxholm Space Rat Atlas version 4.0 to the Brainglobe repository. Using the Brainglobe Atlas API, Brainways can be used to manually register brain slices to any of the Brainglobe-available brain atlases, such as mouse, zebra fish, and humans. Brainways currently provides automatic registration for the Waxholm Space Rat atlas and the Allen mouse brain atlas.

The automatic registration algorithm was trained using 1,970 slices from 5 experiments performed by our group. As some slices had multiple markers, all available markers were used to train the network, for a total of 2,697 images used in the training procedure^2^. To achieve better registration accuracy using limited amounts of data, the algorithm first trained on 40,000 synthetic slices generated by virtually slicing the Waxholm Space rat atlas and the Allen mouse atlas. Each synthetic slice was taken from a random location in the atlas with random 3D rotation and then augmented in various ways (Supplementary 2). The deep neural network architecture of the registration algorithm is based on the Resnet 50 (He et al., 2016) image classification architecture, pretrained on ImageNet (Deng et al., 2009). Three classification heads were added to the baseline architecture, one for AP axis registration, one for visible hemisphere estimation and one for confidence estimation (Fig. 3). Confidence estimation is achieved by training the network to predict whether the AP axis output by the network matches the correct registration. Confidence estimation works well to identify slices that are extremely bright or extremely dark, images with missing tissue, or extremely deformed tissue (Supplementary 2). The rat and mouse models were trained and evaluated separately.

To measure model performance, the AP value output from the network was compared with the AP value annotated by the expert annotators on the test data set. If the AP value of the network matched the value of the annotator by up to 20 voxel units, the registration was considered correct. The rat model achieved 80% accuracy according to this metric in the test data set, with a mean absolute error of 13.93 voxels. The mouse model achieved 90% accuracy, with a mean absolute error of 8.52 voxels. When taking into consideration only slices on which the model was highly confident based on the confidence estimation output, the precision of the rat model increased to 85%. The accuracy of the hemisphere estimation output was 88.7% for the rat model, and was not trained for the mouse model due to the lack of hemisphere information in the mouse data. We expect the registration accuracy to increase as more training data is annotated by researchers using Brainways.

### Rigid Registration

The main goal of the Rigid Registration module is to roughly align the brain tissue in the input image with the atlas slice that was found in the previous step (Fig. 2B). The Rigid Registration module receives two inputs, the coronal slice image and the matching atlas slice, and outputs four rigid registration parameters: left-right translation, up-down translation, horizontal scale, and vertical scale. The Rigid Registration parameters can be viewed and modified using sliders in the Brainways GUI. The registration is visualized by overlapping the input image with a transparent view of the brain regions defined by the atlas (Fig. 1).

**Figure 1.**
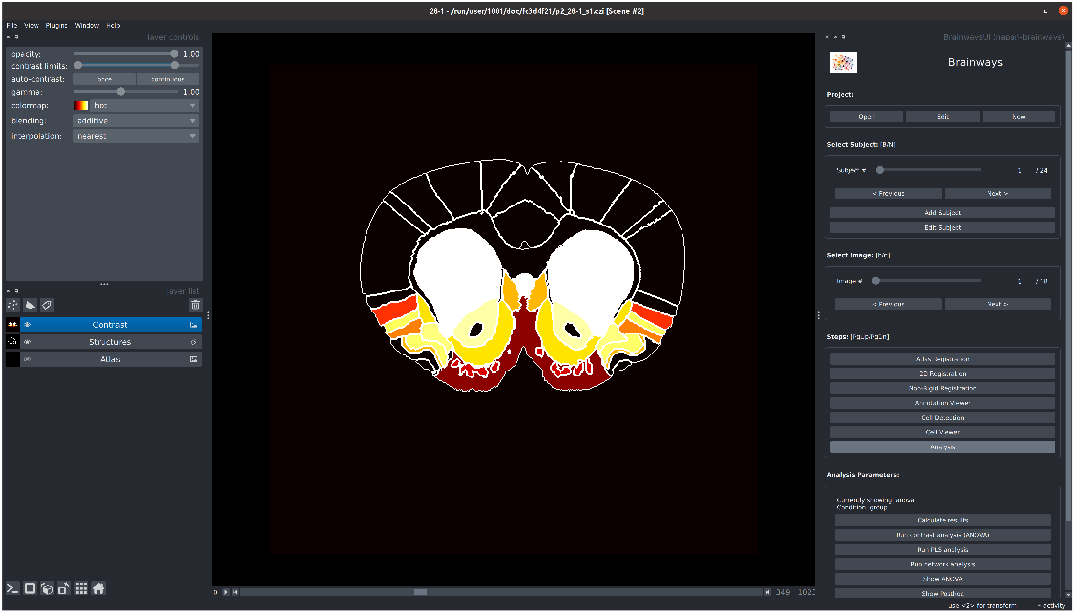
Screenshot taken from the Brainways GUI.

**Figure 2.**
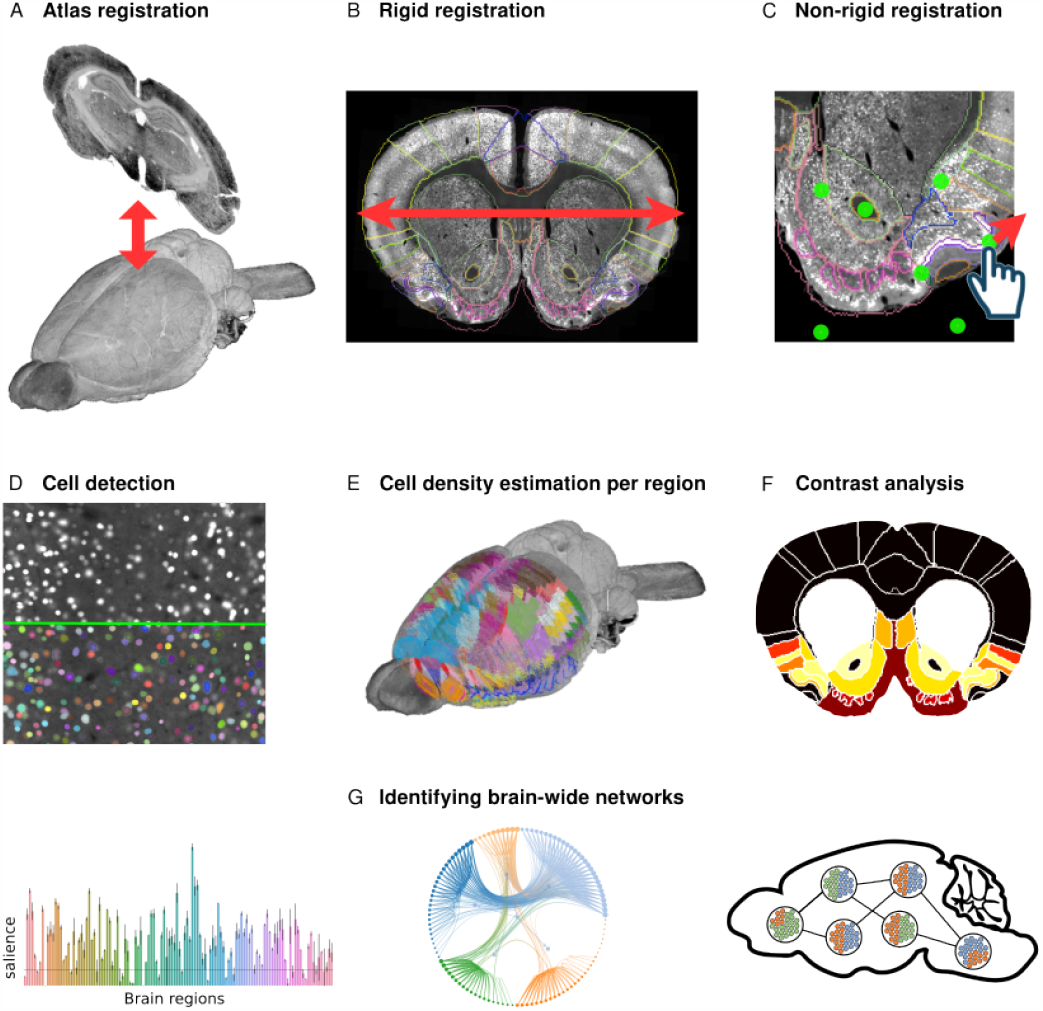
Overview of the Brainways workflow. (**A**) Atlas registration: The images are mapped to the Anterior-Posterior axis of the 3D atlas. (**B**) Rigid registration: The slices are roughly matched to the atlas by moving, scaling, and rotating. (**C**) Non-rigid registration: the slices are elastically deformed to match nonrigid deformations in the slice. (**D**) Cell detection: detection of individual c-Fos activated cells. (**E**) Cell density estimation per region: each detected cell is assigned to its registered region and the number of activated cells in each region is counted. (**F**) A sample of a visual display of the contrast between test conditions provided by Brainways; colors represent the t values. (**G**) Identifying brain-wide networks using Brainways using PLS and network analysis to highlight patterns of regions that drive the contrast between conditions. The network graph is exported by Brainways and visualized via graph-tool Peixoto (2017)

**Figure 3.**
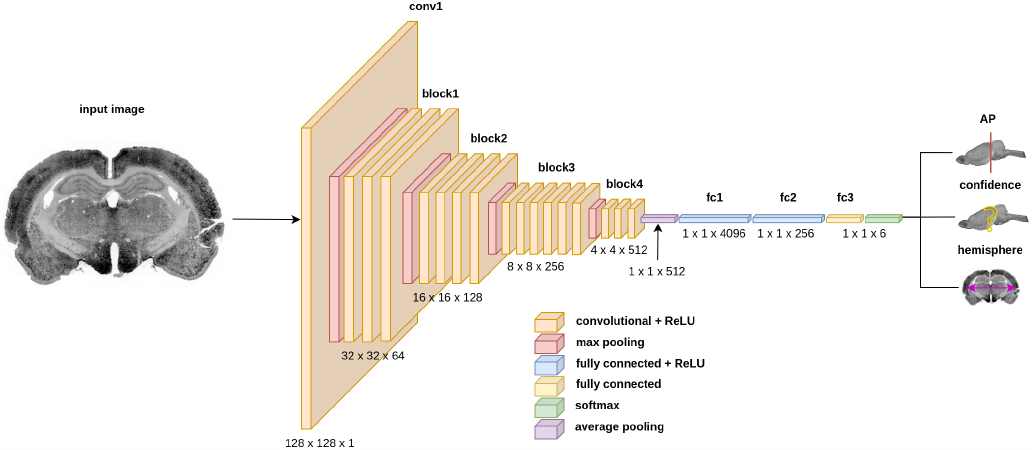
The Brainways algorithm neural network architecture. The architecture is based on Resnet-50 architecture, to which three output heads are appended: 1) Anterior-Posterior location, 2) Confidence, 3) Hemisphere (left/right/both).

Before performing rigid registration between the slice and the atlas, the brain tissue in the image is separated from the background so that only the tissue itself is registered. This is done by binarizing the input image using KMeans on the pixel values in the input image, followed by connected component analysis on binary image (Supplementary 3). Following brain tissue background separation, a bounding box is taken around the brain tissue in the input image and in the atlas slice, and registration parameters are calculated to transform the input box into the atlas box. The automatic Rigid Registration algorithm can be run using a button in the Brainways GUI or via the Python API.

### Non-rigid Registration

Individual histological brain slices often exhibit significant deformations, unlike light-sheet imaging, in which the whole brain is fixed in place. This happens due to the handling and accidental movement of the thin slices, usually between 20-60 *µm*. Experienced experimenters can minimize the extent to which this problem occurs, but it remains a major issue when processing brain slice images. The Non-rigid Registration module receives two inputs, the brain slice image after rigid registration and the matching atlas slice, and outputs keypoints for elastic deformation using Thin Plate Spline (TPS) interpolation. The registration is visualized by overlapping the input image with a transparent view of the brain regions as defined by the atlas.

To achieve elastic deformation, 24 initial *source* points were evenly spaced on the brain tissue and 24 *destination* points were placed at the same locations as the *source* points. Destination points can be moved around using the Brainways GUI, resulting in elastic deformation of the brain tissue that will match the source points to the destination points (Fig. 2C). If the initial 24 point pairs are not sufficient to register key brain regions of interest, additional points can be added or subtracted using the Brainways GUI.

Automatic Non-rigid Registration is supported using the TPS interpolation registration from the Elastix (Klein et al., 2010; Shamonin, 2013) Python package. While Elastix tends to perform good automatic registration to match the outlines of the brain slice to the outlines of the atlas slice, it often fails for more subtle internal structures. Thus, Brainways is designed for experimenters to review the registrations and manually correct them when necessary. In future versions of Brainways, we plan to train a deep neural network to perform the TPS interpolation. Given enough training data, we expect a deep neural network to perform better (and faster) than the current implementation.

### Cell Detection

After registration, the number of positively stained cells in each region is quantified. The Cell Detection module receives the brain slice images and outputs the x- and y-coordinates and the matching brain region for each positively stained cell/nuclei. This is done using an automatic cell detection algorithm, followed by registering the detected cells to the atlas using the registration parameters found in the previous modules.

The StarDist (Schmidt et al., 2018) cell detection algorithm was used for the automatic detection of positively stained cells. StarDist outputs a pixel mask for each detected cell (Fig. 2D), from which several parameters are extracted, such as the central x- and y-coordinates, the area of the cell in microns, and the mean intensity value of each detected cell. These parameters are used for subsequent filtering of false positives. After filtering, each x- and y-coordinates are transformed from the 2D image space to the 3D atlas space by applying the transformations computed by the previous modules. Finally, the corresponding brain region of each cell is computed from its location in the atlas space coordinates. As automatic cell detection for a whole experiment can take many hours, detections can be previewed in the Brainways GUI for detection parameter adjustment before detecting a whole experiment. The process can also be run through the Python API. Brainways supports importing results from external cell detection algorithms via the Import Cell Detections feature.

### Analysis

Following registration and cell counting, the results of the whole experiment (multiple subjects belonging to a single project) are aggregated for subsequent statistical analysis. For each animal and brain region, the total area of the region in microns, the cell count, and the normalized cell count (cell count per 250*µm*^2^) are extracted. Normalized cell count is a better ground for comparison between animals and experiments, because it is independent of variations in region areas between subjects. Detected cells from the whole brain can be viewed in 3D using the Cell Viewer module in the Brainways GUI (Fig. 2E), and can be exported to an Excel file for analysis using external statistical software. The following analyses can be performed from within Brainways on the normalized per-region cell counts.

#### ANOVA contrast analysis

ANOVA contrast analysis can be performed through Brainways to identify and visualize the brain regions that contributed to the contrast between different experimental conditions (Figure 2F). ANOVA is performed on each brain region separately, followed by FDR correction for multiple comparisons (Benjamini et al., 2006). Regions that significantly contributed to the contrast are subjected to post hoc analysis to identify the particular differences between condition pairs.

#### PLS contrast analysis

Task PLS is a multivariate statistical technique that is used to identify optimal neural activity patterns that differentiate between experimental conditions (Mcintosh, 1999; McIntosh et al., 1996). PLS produces a set of mutually orthogonal pairs of latent variables (LV). One element of the LV depicts the contrast, which reflects a commonality or difference between conditions. The other element of the LV, the relative contribution of each brain region (termed here ‘salience’), identifies brain regions that show the activation profile across tasks, indicating which brain areas are maximally expressed in a particular LV. Statistical assessment of PLS is performed using permutation testing for LVs and bootstrap estimation of standard error for the brain region saliences. The significance of latent variables is assessed by permutation testing. The reliability of the salience of the brain region is assessed using bootstrap estimation of standard error. Brain regions with a bootstrap ratio >2.57 (roughly corresponding to a confidence interval of 99%) are considered to be reliably contributing to the pattern. Missing values are interpolated by the average for the test condition.

The results of the analysis are exported to an Excel file and to PNG files containing the salience plot, the LV p values and the LV contrast direction (Figure 5C,D). Brainways uses the *pyls* Python package (Markello, 2023) to perform the PLS analysis.

#### Network graph analysis

To examine the functional connectivity between the different brain regions, Brainways can be used to create a network graph based on inter-region positive cell count correlation matrices. Network nodes consist of the different brain regions as defined in the atlas, and the edges of the network consist of significant correlations between the regions (the significance threshold can be adjusted by the user, by default p<0.05), based on Pearson’s pairwise correlation. The values are FDR corrected for multiple comparisons. The network graph is exported to a graphml file, and can be used with any graph analysis tools and algorithms for further analysis. See the Case Study section for an example analysis using the network graphs.

### Case Study

To investigate the validity of Brainways and demonstrate the advantage of using it over manual quantification methods, we present a Brainways-assisted quantification of a data set from our group (Ben-Ami Bartal et al., 2021). The data set shows images of coronal slices stained for the immediate early gene c-Fos, taken from rats tested in the Helping Behavior Test (HBT, Fig. 4A). The HBT paradigm reflects prosocial motivation in rats who can release a trapped conspecific, thus improving their well-being (Bartal et al., 2011). In the original study, rats selectively helped ingroup members (same-strain conspecifics), but not outgroup members (strangers of an unfamiliar strain, Fig. 4B), in line with previous findings (Ben-Ami Bartal et al., 2014). This data set was originally manually quantified via visual identification of 84 ROIs, followed by ImageJ (Schindelin et al., 2012) intensity-based cell detection for region-by-region quantification (Fig. 4C-D).

**Figure 4.**
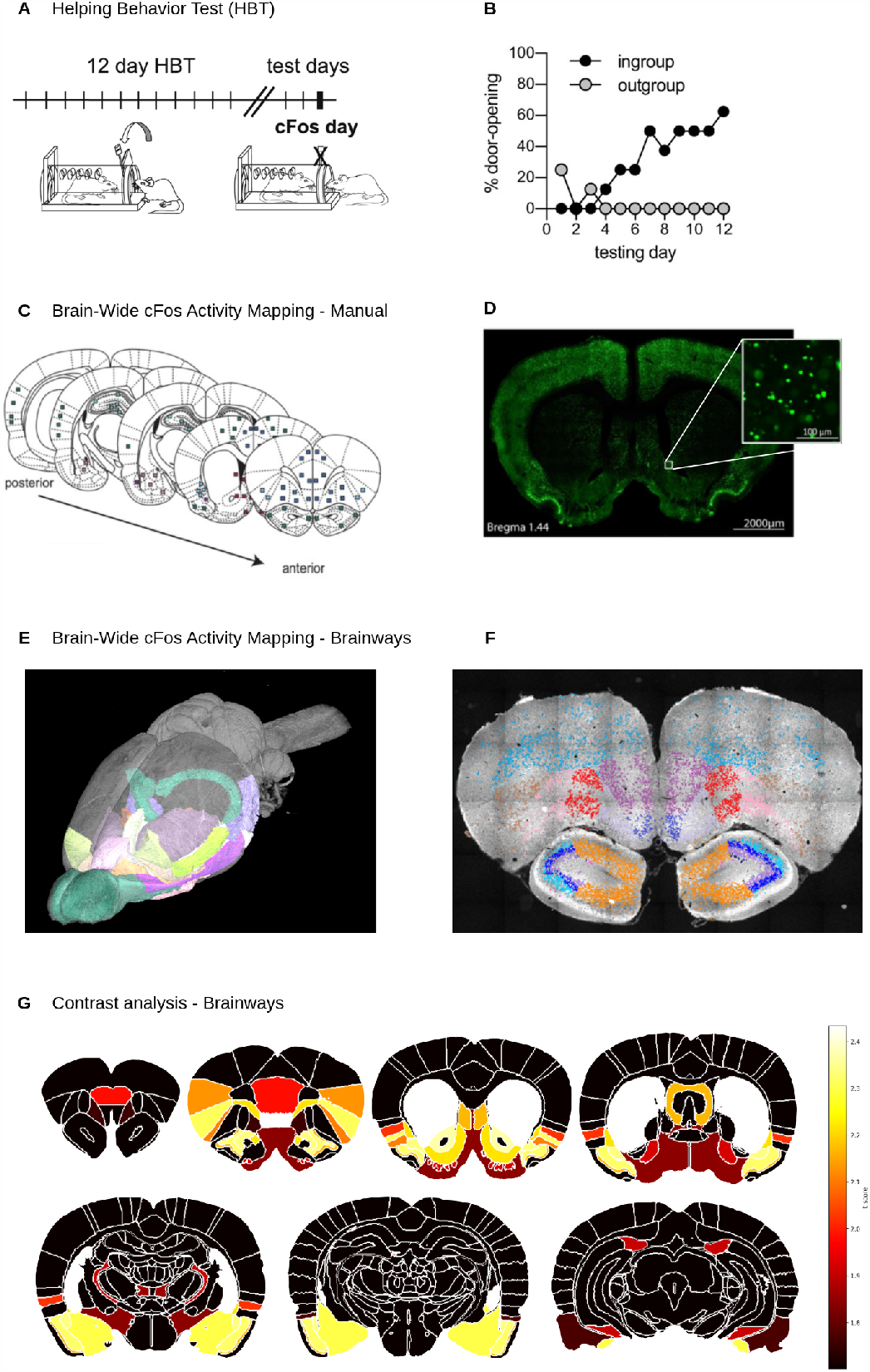
(**A**) Diagram of the Helping Behavior Test. (**B**) Rats released ingroup members but not outgroup members, measured by door-opening over the test days. (**C-D**) Diagram of manual acquisition of c-Fos activity mapping, as was performed in Ben-Ami Bartal et al. (2021). The slices were manually matched to the Waxholm Space Rat Atlas (Papp et al., 2014). From these slices, 84 regions of interest were manually extracted followed by counting c-Fos activated cells in each region. (**E-F**) Brain-wide c-Fos activity mapping as automatically acquired in the Brainways pipeline. All slices were automatically registered to a 3D version of the Waxholm Space Rat Atlas (Osen et al., 2019) to obtain an activity mapping with much higher spatial resolution compared to the previous method. (**G**) ANOVA, post hoc contrast analysis between experimental conditions as visualized using Brainways.

Although this strategy has proven helpful in elucidating the neural mechanisms of prosocial motivation, the analysis was very labor intensive and used only a small fraction of the total available imaged brain tissue. Brainways was used below to demonstrate how these problems can be solved. Brainways analysis resulted in a brain-wide quantification of c-Fos+ cell numbers and was used to re-run the comparison of neural activity patterns between rats under ingroup and outgroup conditions (Fig. 4E-F, see methods). Overall, the resulting reanalysis was highly similar to the original one, providing evidence for the validity of this automated approach, as will be detailed below.

To identify patterns of neural activity associated with each condition, a similar strategy was followed as in the original publication (Ben-Ami Bartal et al., 2021). The neural activity of rats tested in HBT with members of the trapped ingroup or outgroup members was compared with the rats in the control conditions using multivariate task partial least squares analysis (PLS, see Methods, (Mcintosh, 1999; McIntosh et al., 1996)). The original analysis revealed a distinct neural network activated in the presence of a trapped ingroup member during the HBT (Fig. 5A-B). This network included the Insula, ACC, orbitofrontal cortex, and sensory cortices. Importantly, activity in regions associated with the neural reward network, including NAC, LS, VDB, and parts of OFC and PRL were specifically correlated with prosocial motivation and dissociated from arousal around a non-helped trapped outgroup member (Ben-Ami Bartal et al., 2021; Breton et al., 2022). Upon re-acquisition and analysis using Brainways, a similar significant latent variable (LV, p<0.001) emerged, and following boot-strapping and permutations provided a measure of the contribution of each brain region to the significant LV (termed ‘salience’). This analysis revealed an increase in brain-wide activity for rats tested in HBT with trapped ingroup or outgroup members compared to rats in the baseline condition (Fig. 5C, black bars). This network included regions in the sensory cortex, frontal regions, limbic and reward regions (Fig. 5C, colored bars). Notably, as previously reported, these regions include areas that have been shown to participate in empathy in humans (Shamay-Tsoory and Lamm, 2018; Decety et al., 2016) and rodents (Meyza and Knapska, 2018; Wu and Hong, 2022; Walsh et al., 2023; Paradiso et al., 2021). A direct comparison between the ingroup and outgroup conditions also produced a significant LV, (LV1, p<0.01), and pointed to a distinct network that was significantly more active for ingroup members compared to outgroup members (Fig. 5D).

**Figure 5.**
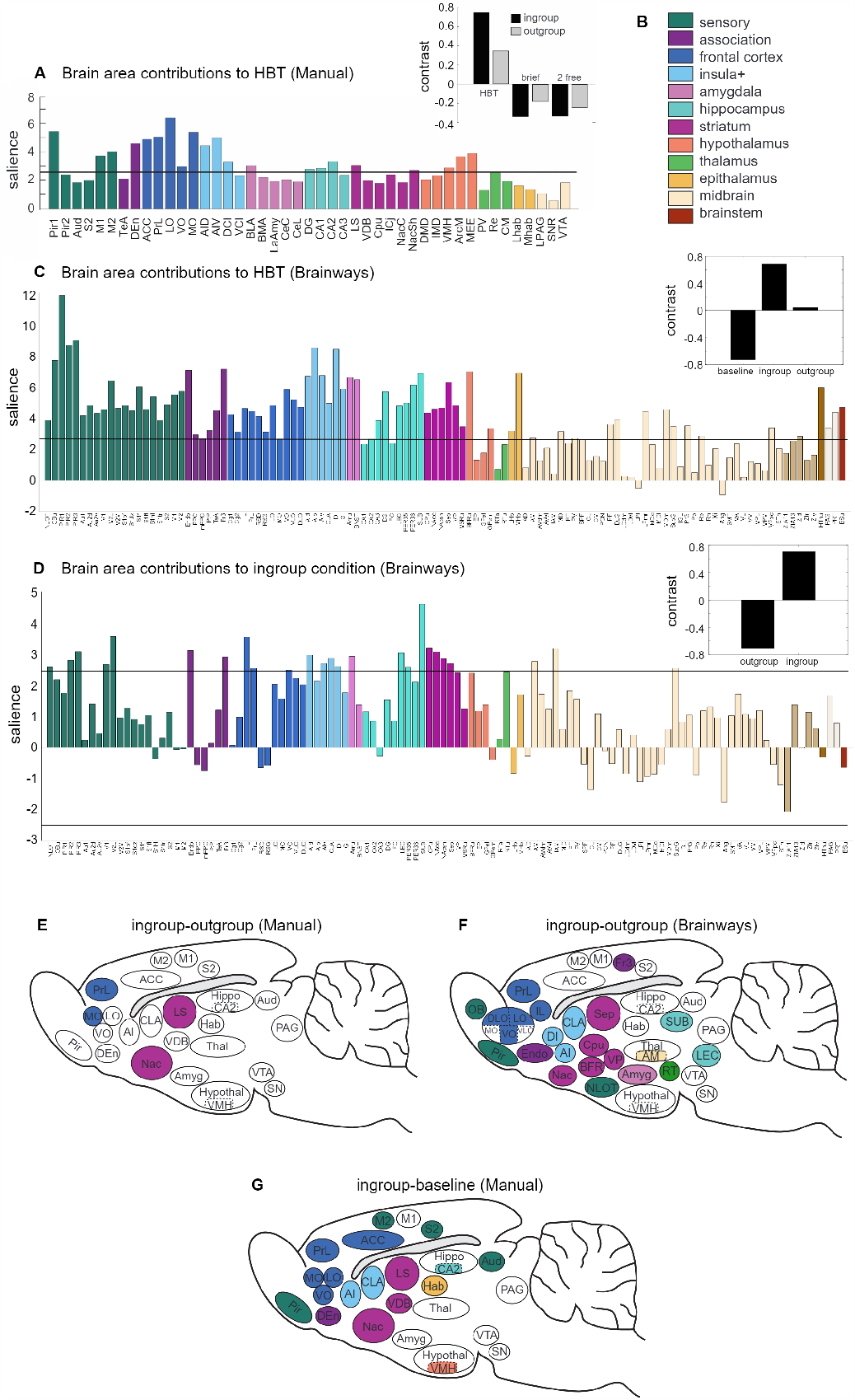
(**A**) Partial least square (PLS) task analysis reveal a neural activity pattern unique to the HBT task (based on the manual acquisition, taken from BenAmi Bartal et al. (2021)). Salience represents the z score of boot-strapping tests, with regions that cross the black threshold lines significantly contributing to the contrast depicted in the inset (p<0.01, black bars). (**B**) Color-coded legend of brain regions (**C**) Brainways-assisted acquisition of the same data reveals a more detailed and significant neural activity pattern unique to the HBT task. (**D**) Contrasting the ingroup and outgroup conditions reveal a neural activity pattern unique to the ingroup condition. (**E**) Diagram of a rat brain showing regions significantly active (in color) for the HBT ingroup condition, derived using manual data acquisition, (**F**) or using Brainways-assisted acquisition. (**G**) The contrast comparing the ingroup and outgroup conditions using Brainways resembles the contrast comparing the ingroup and baseline conditions using manual acquisition. The complete abbreviation table is found in Supplementary 1.

Next, the number of c-Fos+ cells was compared for the ingroup, outgroup, and baseline conditions (ANOVA, FDR corrected for multiple comparisons (Benjamini et al., 2006)). The results of this analysis expand on the original publication where only four regions (MO, PrL, NAC, LS) reached the significance threshold for this comparison (Fig. 5E). With the exception of the MO, the three other regions (NAC, Septum, PrL) were replicated. Furthermore, areas that were not previously quantified also proved significant, including the IL, VP, and parts of the thalamus (LEC, Subiculum, Fig. 5F). Multiple regions that previously emerged as active in the HBT ingroup condition compared to baseline were replicated (Fig. 5G), including the claustrum, insula, piriform and endopiriform cortex, basal forebrain, Cpu, and the amygdala. Thus, the Brainways-assisted reanalysis reproduced the original findings and revealed the expanded empathic helping network more fully, providing further evidence in the role of this network in prosocial motivation.

### Network Analysis

To examine the functional connectivity between the different brain regions, network graphs based on inter-regional correlation matrices of c-Fos+ cell numbers were generated through Brainways based on a Pearson pairwise correlation matrix. The resulting graph was clustered to identify community structures and visualized using Cytoscape (Shannon et al., 2003) (Fig. 6). The resulting network graph shows a central cluster that includes multiple regions of the prosocial network (NAc, MO, Insula, IL, claustrum, and auditory sensory regions). A second cluster consisted of frontal regions (RS, ACC, PrL, OFC, Fr3) and multiple regions in the visual and somatosensory cortex. This cluster also includes the hippocamapal regions (CA2, CA1, DLG) and the lateral habenula. The third and fourth clusters were mostly made up of thalamic subregions. This network, based on the Brainways quantification, indicates that the regions involved in prosocial motivation make up a distributed brain-wide network connecting affective and motivational regions that are functionally connected during HBT. (Ben-Ami Bartal et al., 2021)..

**Figure 6.**
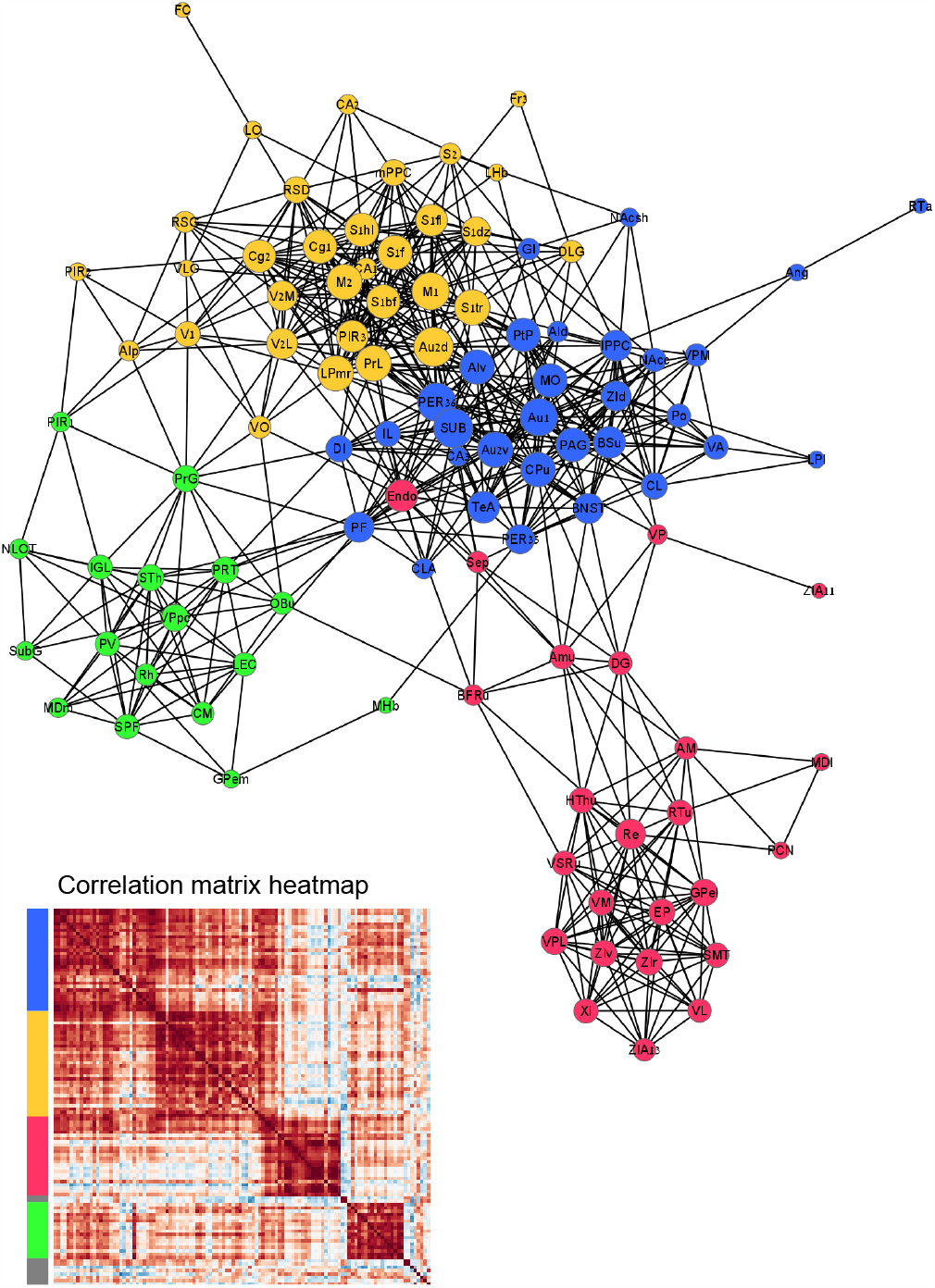
Network graph and corresponding interregion correlation matrix heatmap representing the functional connectivity between different brain regions in the ingroup HBT condition.

## Discussion

Here we offer an open source tool for automatic registration, quantification, and analysis of stained coronal slices. Moreover, Brainways aims to provide a synthesis of the brain-wide network associated with a manipulation of interest. Above we demonstrate the utility of Brainways in outlining the neural connectome of prosocial motivation, and validate the accuracy of Brainways against manual quantification of the same data. Brainways is a useful tool for researchers working with fluorescence tagging, who are particularly interested in visualizing the brain-wide network associated with their marker of interest. Although Brainways was conceptualized for analysis of neural activity markers (EIG like Fos), other use-cases include receptor mapping, population identification, and via co-labeling, activity-identity mapping (e.g. receptors on active cells).

Brainways is optimized for analysis of coronal slices, the most commonly used brain imaging method in animal studies across model organisms. Brainways is an easy-to-use tool allowing fast and global quantification of large amounts of data, especially designed for laboratories lacking skilled programmers. Brainways allows for the automatic registration of coronal slices to a 3D digital atlas, supports elastic deformations, stained cell detection, and aggregating cell counts for each brain region and animal in an experiment. Brainways provides integrated ANOVA, PLS and graph network analyses, providing a way to examine the neural patterns of activity associated with different task conditions. This is especially useful for looking at data with relatively low number of rows (i.e. subjects) and a large number of columns (regions) in a non-biased way. This approach leads to identification of central hubs that can be used to identify ROIS for manipulation. Automatic registration is performed using a deep neural network algorithm that receives coronal images as input and outputs the 3D registration parameters.

Brainways was trained on the rat Waxholm and on the Allen mouse atlases and is best optimized for these species. However, any 3D atlas can be used by novel users, who may perform manual registration. Our aim is to train more model species by working with the community according to their needs. Brainways uses the Brainglobe API as its backbone for accessing the 3D atlas, which makes Brainways compatible with all atlases offered by Brainglobe, including mice, zebrafish, and human atlases.

To encourage further development in the field, the Brainways code is publicly shared as open-source software ^3^. The software can be used by researchers with no programming skills through a GUI, or via a Python API for users who wish to utilize Brainways as part of other scripts or programs.

To demonstrate the utility of Brainways, we have reanalyzed a previously published experiment that was originally analyzed using manual acquisition methods. Brainways-assisted analysis largely replicated the neural activity pattern found in the previous study and revealed additional regions that were not included in the manual quantification. Using Brainways, the amount of tissue covered in the analysis increased by over a hundred times, which contributed to greater statistical significance of the results by increasing the signal strength compared to the previous study. The similarity of the findings is encouraging, both strengthening the validity of the identified ROIs and the reliability of the output provided by Brainways. Notably, even though the analysis was based on the same data set, closely replicating the previous results is not a trivial outcome, because the reanalysis covered a significantly larger surface area, and was processed from the raw images by different experimenters who did not see the original data.

## Limitations

The current approach is highly beneficial for rapidly gaining brain-wide quantification of fluorescence. However, there are some caveats that should be considered. The resolution provided by Brainways suffers on two levels. One is the number of coronal slices provided by the user. As sampling increases, so does the resolution of the resulting network. However, since using Brainways significantly shortens the time from scanning the slides to analysis, it becomes much more feasible to process a higher number of slices per brain.

Another limit is the level of parcellation offered by the atlas in use. Here, for rats, the digital Waxholm Space Rat Atlas was used, and some key regions, such as the amygdala, hypothalamus, and basal forebrain, are not subdivided any further. Encouragingly, this atlas is under active development and we hope that the next version will include these important subregions. Furthermore, this is not an issue for investigators working with mice, as the existing Allen mouse brain atlas is highly parcellated.

Another limitation of Brainways is that it tends to underfit small parcellated subregions in the Non-rigid Registration phase. Therefore, users are advised to manually scan and fix fit errors during this phase, especially if they focus on differences between specific subregions of interest. We plan to train a non-rigid registration algorithm based on the data collected using the Brainways GUI in order to alleviate this issue. It is important to note that regardless of model accuracy, attentive processing of the sections is an inherent part of ensuring data quality in any software solution. Brainways GUI is especially designed to facilitate and speed up the visual scanning process and manual adjustment of the automatic registrations.

Finally, Brainways currently processes a single channel automatically. Yet, co-labeling of several fluorescent markers is often utilized. Brainways supports importing cell detections of multiple channels (e.g. from QuPath), registering them to the atlas, and exporting a co-labeling cell count for each brain region and each animal in the experiment. Brainways currently only supports cell detection of a single stain from within the software, and we plan to include cell detection of multiple stains in the near future.

In conclusion, Brainways is a novel open source AI-based software solution that can lead to novel discoveries of neurobiological mechanisms through its ability to provide quick and accurate automated network analysis from a data set of stained coronal slices. Brainways is offered freely to the community in the hope that it will lead to a leap forward in the field and promote collaborative efforts for meta-analyses revealing basic neural function.

## Methods

The methods used to obtain the c-Fos data used in the case study have been described in detail in BenAmi Bartal et al. (2021). Thus, they are briefly described below.

### Dataset curation

To perform the registrations for the reanalysis, four annotators blind to task condition each received a portion of the original data set. Annotators used Brainways GUI to perform automatic registrations for each slice, followed by manual fit adjustments when they perceived a mismatch between the atlas and the digital scan. The complete data set was then reviewed by a fifth annotator to create the final registrations.

### Experimental design

#### Animals

Rat studies were performed according to protocols approved by the Institutional Animal Care and Use Committee of the University of California, Berkeley. Rats were kept on a 12-h light-dark cycle and received food and water ad libitum. In total, 83 rats were tested across all experiments. Adult pair-housed male SD rats (‘SD,’ age p60–p90 days) were used as free and trapped ingroup rats (Charles River, Portage, MI). Adult male Long–Evans rats were used as trapped outgroup rats (LE, Envigo, CA). All rats ordered were allowed a minimum of 5 days to acclimate to the facility prior to beginning testing.

#### Helping Behavior Test

The HBT was performed as previously described (Bartal et al., 2011). During the HBT, a free rat was placed in an open arena containing a rat trapped inside a restrainer. The free rat could help the trapped rat by opening the restrainer door with its snout, thereby releasing the trapped rat. One-hour sessions were repeated daily over a 2-week period. At 40 min, the door was opened half-way by the experimenter; this was typically followed by the exit of the trapped rat and was aimed at preventing learned helplessness. Door opening was counted as such when performed by the free rat before the half-way opening point. Rats that learned to open the restrainer and consistently opened it on the last three days of testing were labeled ‘openers’. On the last day of testing, the restrainer was latched shut throughout the 60 min session and rats were perfused immediately following behavioral testing.

#### Immunohistochemistry

On the last day of testing, the animals were sacrificed within 90 minutes from the beginning of the session, at the peak expression of the early immediate gene product c-Fos. Brains were extracted following perfusion, frozen, sliced at 40 *µ*m and stained for c-Fos. The sections were stained for c-Fos using rabbit anti-c-Fos antiserum (ABE457; Millipore, 1:1000; 1% NDS; 0.3% TxTBS) with Alexa Fluor 488 conjugated donkey anti-rabbit antiserum (AF488; Jackson, 1:500; 1% NDS; 0.3% TxTBS). The sections were further stained in DAPI (1:40,000). Details of the imaging and manual acquisition methods can be found in the original publications (Ben-Ami Bartal et al., 2021). Immunostained tissue was imaged at 10× using a wide-field fluorescence microscope (Zeiss AxioScan) and processed in Zen software.

### Statistical analysis

Statistic methods for PLS analysis, network graph and ANOVAs are outlined in the Results section detailing the functionality of the Analysis module.

## Acknowledgements

The authors thank the following people for their help with this manuscript: Keren Ruzal, Estherina Trachtenberg, Einat Bigelman, and Reut Hazani. This work was supported by the Azrieli Foundation, the Israel Science Foundation (ISF), and the Tel Aviv University Center for AI and Data Science (TAD).

## Footnotes

1 Brainways: https://github.com/bkntr/brainways; Brainways GUI: https://github.com/bkntr/napari-brainways; Brainways registration model: model: https://github.com/bkntr/brainways-reg-model

2 All our data is publicly available, more details in the GitHub page.

3 Brainways: https://github.com/bkntr/brainways; Brain-ways GUI: https://github.com/bkntr/napari-brainways; Brainways registration https://github.com/bkntr/brainways-reg-model

