## Supplemental File for "Brainways: An AI-based Tool for Automated Registration, Quantification and Generation of Brain-wide Activity Networks Based on Fluorescence in Coronal Slices"

### Supplementary Information

#### Supplementary Note 1: Brain region abbreviation table

| acronym | name |
|---|---|
| AD | Anterodorsal thalamic nucleus |
| AId | Agranular insular cortex dorsal area |
| AIp | "Agranular insular cortex, posterior area " |
| AIv | "Agranular insular cortex, ventral area" |
| AM | Anteromedial thalamic nucleus |
| Amu | "Amygdaloid area, unspecified" |
| Ang | Angular thalamic nucleus |
| Au1 | Primary auditory area |
| Au2d | "Secondary auditory area, dorsal part" |
| Au2v | "Secondary auditory area, ventral part" |
| AVdm | "Anteroventral thalamic nucleus, dorsomedial part" |
| AVvl | "Anteroventral thalamic nucleus, ventrolateral part" |
| BFRu | "Basal forebrain region, unspecified" |
| BNST | Bed nucleus of the stria terminalis |
| BSu | "Brainstem, unspecified" |
| CA1 | Cornu ammonis 1 |
| CA2 | Cornu ammonis 2 |
| CA3 | Cornu ammonis 3 |
| Cg1 | Cingulate area 1 |
| Cg2 | Cingulate area 2 |
| CL | Central lateral thalamic nucleus |
| CLA | Claustrum |
| CM | Central medial thalamic nucleus |
| CNIC | "Inferior colliculus, central nucleus" |
| CPu | Caudate putamen |
| DG | Dentate gyrus |
| DI | Dysgranular insular cortex |
| DLG | Dorsal lateral geniculate nucleus |
| DLO | Dorsolateral orbital area |
| ECIC | "Inferior colliculus, external cortex" |
| Endo | Endopiriform nucleus |
| EP | Entopeduncular nucleus |
| Eth | Ethmoid-Limitans nucleus |
| FC | Fasciola cinereum |
| FoF | Fields of Forel |
| Fr3 | Frontal association area 3 |
| GI | Granular insular cortex |
| GPel | "Globus pallidus external, lateral part" |
| GPem | "Globus pallidus external, medial part" |
| HThu | "Hypothalamic region, unspecified" |
| IAM | Interanteromedial thalamic nucleus |
| IGL | Intergeniculate leaflet |
| IL | Infralimbic area |
| IMD | Intermediodorsal thalamic nucleus |
| IP | Interpeduncular nucleus |
| LDdm | "Laterodorsal thalamic nucleus, dorsomedial part" |
| LDvl | "Laterodorsal thalamic nucleus, ventrolateral part" |
| LEC | Lateral entorhinal cortex |

| acronym | name |
|---|---|
| LHb | Lateral habenular nucleus |
| LO | Lateral orbital area |
| LPl | "Lateral posterior thalamic nucleus, lateral part" |
| LPmc | "Lateral posterior thalamic nucleus, mediocaudal part" |
| LPmr | "Lateral posterior thalamic nucleus, mediorostral part" |
| lPPC | "Parietal association cortex, lateral area" |
| M1 | Primary motor area |
| M2 | Secondary motor area |
| MDc | "Mediodorsal thalamic nucleus, central part" |
| MDl | "Mediodorsal thalamic nucleus, lateral part" |
| MDm | "Mediodorsal thalamic nucleus, medial part" |
| MEC | Medial entorhinal cortex |
| MGd | "Medial geniculate body, dorsal division" |
| MGm | "Medial geniculate body, medial division" |
| MGmz | "Medial geniculate body, marginal zone" |
| MGsg | "Medial geniculate body, suprageniculate nucleus" |
| MGv | "Medial geniculate body, ventral division" |
| MHb | Medial habenular nucleus |
| MO | Medial orbital area |
| mPPC | "Parietal association cortex, medial area" |
| NAcc | "Nucleus accumbens, core" |
| NAcsh | "Nucleus accumbens, shell" |
| NLOT | Nucleus of the lateral olfactory tract |
| OBu | "Olfactory bulb, unspecified" |
| PAG | Periaqueductal gray |
| PaS | Parasubiculum |
| PCN | Paracentral thalamic nucleus |
| PER35 | Perirhinal area 35 |
| PER36 | Perirhinal area 36 |
| PF | Parafascicular thalamic nucleus |
| PIL | Posterior intralaminar nucleus |
| PIR1 | "Piriform cortex, layer 1" |
| PIR2 | "Piriform cortex, layer 2" |
| PIR3 | "Piriform cortex, layer 3" |
| Pn | Pontine nuclei |
| Po | Posterior thalamic nucleus |
| Pot | "Posterior thalamic nuclear group, triangular part" |
| PP | Peripeduncular nucleus |
| PrG | Pregeniculate nucleus |
| PrL | Prelimbic area |
| PrS | Presubiculum |
| PRT | Pretectal region |
| RT | Reticular (pre)thalamic nucleus |
| PT | Parataenial thalamic nucleus |
| PtP | "Parietal association cortex, posterior area " |
| PV | Paraventricular thalamic nuclei (anterior and posterior) |
| Re | Reuniens thalamic nucleus |
| Rh | Rhomboid thalamic nucleus |
| RRe | Retroreuniens thalamic nucleus |
| RSD | Retrosplenial dysgranular area |
| RSG | Retrosplenial granular area |
| RTa | "Reticular (pre)thalamic nucleus, auditory segment" |
| RTu | "Reticular (pre)thalamic nucleus, unspecified" |

| acronym | name |
|---|---|
| S1bf | "Primary somatosensory area, barrel field" |
| S1dz | "Primary somatosensory area, dysgranular zone" |
| S1f | "Primary somatosensory area, face representation" |
| S1fl | "Primary somatosensory area, forelimb representation" |
| S1hl | "Primary somatosensory area, hindlimb representation" |
| S1tr | "Primary somatosensory area, trunk representation" |
| S2 | Secondary somatosensory area |
| Sag | Nucleus sagulum |
| Sep | Septal region |
| SMn | Nucleus of the stria medullaris |
| SMT | Submedius thalamic nucleus |
| SNc | "Substantia nigra, compact part" |
| SNl | "Substantia nigra, lateral part" |
| SNr | "Substantia nigra, reticular part" |
| SPF | Subparafascicular nucleus |
| STh | Subthalamic nucleus |
| SUB | Subiculum |
| SubG | Subgeniculate nucleus |
| SuD | Deeper layers of the superior colliculus |
| SuG | Superficial gray layer of the superior colliculus |
| TeA | Temporal association cortex |
| V1 | Primary visual area |
| V2L | "Secondary visual area, lateral part" |
| V2M | "Secondary visual area, medial part" |
| VA | Ventral anterior thalamic nucleus |
| VL | Ventrolateral thalamic nucleus |
| VLO | Ventrolateral orbital area |
| VM | Ventromedial thalamic nucleus |
| VO | Ventral orbital area |
| VP | Ventral pallidum |
| VPL | Ventral posterolateral thalamic nucleus |
| VPM | Ventral posteromedial thalamic nucleus |
| VPpc | "Ventral posterior nucleus of the thalamus, parvicellular part" |
| VSRu | "Ventral striatal region, unspecified" |
| VTA | Ventral tegmental area |
| Xi | Xiphoid thalamic nucleus |
| ZIA11 | "Zona incerta, A11 dopamine cells" |
| ZIA13 | "Zona incerta, A13 dopamine cells" |
| ZIc | "Zona incerta, caudal part" |
| ZId | "Zona incerta, dorsal part" |
| ZIr | "Zona incerta, rostral part" |
| ZIv | "Zona incerta, ventral part" |

### Supplementary Note 2: The Brainways registration algorithm

The Brainways network takes as input a low-resolution image of a histological brain slice and outputs six parameters: 1) Anterior-Posterior location, 2) Horizontal rotation, 3) Sagittal rotation, 4) Frontal rotation, 5) Hemisphere (left/right/both), and 6) Confidence. We use the standard Resnet 50 (He et al., 2016) architecture pretrained on the ImageNet (Deng et al., 2009) dataset, to which we added 6 classification heads, one for each output parameter. The network is trained in two phases - first, unsupervised training using synthetically generated slices, followed by supervised training using real slices annotated by expert neuroscientists using the Brainways software.

##### Numeric outputs

Following previous work in the field of deep learning, which shows that deep learning models perform better in classification problems over regression problems, we cast the numeric outputs to classification outputs. For each of

the numeric outputs (AP, Horizontal rotation, etc.), instead of directly outputting a number from the model, we split the range of each output $i$ to $n_i$ bins. The minimal and maximal values of that range $min_i, max_i$ are pre-determined based on the training data. The range $(min_i, max_i)$ is then divided into $n_i$ bins, and the model chooses a bin based on the input. The numeric value of the model output is the central value of the chosen bin. Numeric ranges for each output are given in Table S1.
For each numeric output $i$, let $y_i$ be the numeric ground truth of output $i$, let $b_i$ be the bin that corresponds to that value, and let $b'_i$ the predicted bin of the network for output $i$. The loss $L_i$ is calculated as follows:

$$L_i = NLLLoss\left(b_i, b'_i\right)$$

| | # Bins | Min Value | Max Value |
|---|---|---|---|
| AP | 512 | 122 | 799 |
| Hor. Rot. | 7 | -7 | 7 |
| Sag. Rot. | 7 | -7 | 7 |
| Fro. Rot. | 45 | -45 | 45 |

**Table S1.** Number of bins and maximal range of each of the Brainways algorithm numeric outputs.

##### Confidence estimation

The Brainways algorithm is trained to output an estimation of its confidence in the correctness of the automatic registration. This is important because it allows researchers to quickly identify potentially inaccurate registrations, allowing for manual verification and correction if needed. This helps to ensure the reliability and accuracy of the resulting overall registration. It also helps to prioritize the manual registration of certain brain slices, allowing for more efficient use of resources.
The ground-truth label for confidence estimation is calculated in the following way. Let $y_{ap}$ be the AP location of the slice in voxel units, and $y'_{ap}$ the AP location predicted by the model in voxel units. The confidence label $y_{conf}$ is calculated as follows:

$$y_{conf} = \begin{cases} 1, & \text{if } \left|y_{ap} - y'_{ap}\right| < 20 \\ 0, & \text{otherwise} \end{cases}$$

And confidence loss $L_{conf}$ is calculated:

$$L_{conf} = NLLLoss\left(y_{conf}, y'_{conf}\right)$$

##### Loss

The loss of the model is the sum of all output losses and the confidence estimation loss:

$$loss = L_{ap} + L_{hr} + L_{sr} + L_{fr} + L_{hem} + L_{conf}$$

##### Training on synthetic slices

In the Unsupervised Training phase, the network is trained on 20,000 synthetically generated histological brain slices, generated from the 3D WHS SD 39$\mu m$ rat atlas (Osen et al., 2019; Papp et al., 2014) for the rat model, and from the Allen 3D mouse atlas (Wang et al., 2020) for the mice model, both accessed using the BrainGlobe API (Claudi et al., 2020). Each slice was taken from a random location in the anterior-posterior axis, with random 3D rotation in the frontal, horizontal and sagittal axes. Each slice was subjected to the following augmentation procedures:

1. Randomly crop a single hemisphere.
2. Crop image to contain only non-background pixels.
3. Randomly adjust contrast.
4. Resize image to $W \times H$.
5. Random lighten dark areas.
6. Random zero darker areas to simulate tissue tear.
7. Add random light patches.
8. Random affine projection.
9. Random elastic deformation.

##### Fine-tuning on real data

After training on synthetic slices, the network is then fine-tuned on real images of brain slices, annotated using the Brainways pipeline.

***Data - rats.*** Whole brains from three different experiments were annotated for training and validating the Brainways algorithm for rats. One experiment was allocated exclusively for the test phase, and the other two experiments were split 80% for training and 20% for validation. For the training of the model, all image channels were used (DAPI, cFos and Retrograde, where available). For validation and testing, only the cFos channel was used (Table S2).

| | # Images | # Unique Images | # Experiments |
|---|---|---|---|
| Train | 1444 | 717 | 2 |
| Validation | 178 | 178 | 2 |
| Test | 907 | 907 | 1 |

**Table S2.** Brainways algorithm data counts - rats.

***Data - mice.*** Whole brains from a single experiment were annotated for training and validating the Brainways algorithm for mice. The images from the experiment were split 50% for training, 25% for validation and 25% for test (Table S3).

| | # Images | # Unique Images | # Experiments |
|---|---|---|---|
| Train | 84 | 84 | 1 |
| Validation | 42 | 42 | 1 |
| Test | 42 | 42 | 1 |

**Table S3.** Brainways algorithm data counts - mice.

#### Supplementary Note 3: Tissue Background Separation Algorithm

The tissue-background separation algorithm is described below:

1. Get image $I$ with $p$ pixels.
2. Clip pixel values of $I$ to be between 0th and 50th percentiles of all pixel values.
3. Run KMeans with two clusters on all image pixel values to get a quantized binary image $Q$.
4. Find connected components $C_i$ in $Q$
5. For each connected component $C_i$, count the number of pixels in the component to get $N_i$.
6. Find the largest connected component $L = \arg\max\{N_i\}$.
7. Remove from $Q$ very small components, such that $N_i \leq 0.01 \cdot p$
8. Remove from $Q$ small components around the edges of the image, such that $C_i$ touches the edge of the image and $N_i \leq 0.5 \cdot N_L$.
9. Return $Q$.

#### Bibliography (Supplementary)


Claudi, F., Petrucco, L., Tyson, A., Branco, T., Margrie, T., and Portugues, R. (2020). BrainGlobe Atlas API: a common interface for neuroanatomical atlases. *Journal of Open Source Software*, 5 (54):2668. doi: 10.21105/joss.02668.

Deng, J., Dong, W., Socher, R., Li, L.-J., Kai Li, and Li Fei-Fei. ImageNet: A large-scale hierarchical image database. In *2009 IEEE Conference on Computer Vision and Pattern Recognition*, pages 248–255, Miami, FL, (2009). IEEE. ISBN 978-1-4244-3992-8. doi: 10.1109/CVPR.2009.5206848. URL https://ieeexplore.ieee.org/document/5206848/.

He, K., Zhang, X., Ren, S., and Sun, J. Deep Residual Learning for Image Recognition. In *2016 IEEE Conference on Computer Vision and Pattern Recognition (CVPR)*, pages 770–778, Las Vegas, NV, USA, (2016). IEEE. ISBN 978-1-4673-8851-1. doi: 10.1109/CVPR.2016.90. URL http://ieeexplore.ieee.org/document/7780459/.

Osen, K. K., Imad, J., Wennberg, A. E., Papp, E. A., and Leergaard, T. B. (2019). Waxholm Space atlas of the rat brain auditory system: Three-dimensional delineations based on structural and diffusion tensor magnetic resonance imaging. *NeuroImage*, 199:38–56. doi: 10.1016/j.neuroimage.2019.05.016.

Papp, E. A., Leergaard, T. B., Calabrese, E., Johnson, G. A., and Bjaalie, J. G. (2014). Waxholm Space atlas of the Sprague Dawley rat brain. *NeuroImage*, 97:374–386. doi: 10.1016/j.neuroimage.2014.04.001.

Wang, Q., Ding, S.-L., Li, Y., Royall, J., Feng, D., Lesnar, P., Graddis, N., Naeemi, M., Facer, B., Ho, A., Dolbeare, T., Blanchard, B., Dee, N., Wakeman, W., Hirokawa, K. E., Szafer, A., Sunkin, S. M., Oh, S. W., Bernard, A., Phillips, J. W., Hawrylycz, M., Koch, C., Zeng, H., Harris, J. A., and Ng, L. (2020). The Allen Mouse Brain Common Coordinate Framework: A 3D Reference Atlas. *Cell*, 181(4):936–953.e20. doi: 10.1016/j.cell.2020.04.007.
